# Forest *Saccharomyces paradoxus* are robust to seasonal biotic and abiotic changes

**DOI:** 10.1101/2020.04.27.063289

**Authors:** Primrose J. Boynton, Dominika Wloch-Salamon, Doreen Landermann, Eva H. Stukenbrock

## Abstract

Microorganisms are famous for adapting quickly to new environments. However, most evidence for rapid microbial adaptation comes from laboratory experiments or domesticated environments, and it is unclear how rates of adaptation scale from human-influenced environments to the great diversity of wild microorganisms. We examined potential monthly-scale selective pressures in the model forest yeast *Saccharomyces paradoxus*. Contrary to expectations of seasonal adaptation, the *S. paradoxus* population was stable over four seasons in the face of abiotic and biotic environmental changes. While the *S. paradoxus* population was diverse, including 41 unique genotypes among 192 sampled isolates, there was no correlation between *S. paradoxus* genotypes and seasonal environments. Consistent with observations from other *S. paradoxus* populations, the forest population was highly clonal and inbred. This lack of recombination, paired with population stability, implies that *S. paradoxus* evolved the phenotypic plasticity needed to resist seasonal environmental fluctuations long ago, and that individual *S. paradoxus* are generalists with regard to seasonal environments. Similarly, while the forest population included diversity among phenotypes related to intraspecific interference competition, there was no evidence for active coevolution among these phenotypes. At least ten percent of the forest *S. paradoxus* individuals produced “killer toxins”, which kill sensitive *Saccharomyces* cells, but the presence of a toxin-producing isolate did not predict resistance to the toxin among nearby isolates. How forest yeasts acclimate to changing environments remains an open question, and future studies should investigate the physiological responses that allow microbial cells to cope with environmental fluctuations in their native habitats.

## Introduction

All organisms, large and small, live in changing environments. These changing environments are diverse: environmental changes have frequencies ranging from fast to slow; are directional, random, or cyclic; and come from abiotic or biotic sources (Andrews & Brasier, 2005; Kidwell, 2015; Meyers & Bull, 2002). Organisms’ responses to changes are equally diverse: they can evolve to adapt to new environments; acclimate through temporary changes in physiology or behavior; adjust their ranges; or respond through a combination of these and other processes (Nadeau & Urban, 2019). *Microbial* responses to changing environments are emerging as an important theme of current research, especially as researchers grapple with human impacts on climate and biodiversity (Antwis et al., 2017). Responses to climate change among more easily studied macroorganisms are varied and difficult to extend to microorganisms (Rowe et al., 2015; A. B. Smith et al., 2019). How rapid evolutionary adaptation helps microbial populations respond to rapid environmental changes is an open question in microbial ecology.

Most information about microbial responses to environmental changes comes from human influenced environments, in which rapid adaptation is common. Rapid adaptation is frequently observed in laboratory experimental evolution: under controlled laboratory conditions, microbial fitness in new environments increases substantially over timescales of days to months (Gómez & Buckling, 2011; Luria & Delbrück, 1943; Rafaluk-Mohr, Ashby, Dahan, & King, 2018; Rainey & Travisano, 1998). For example, populations of the model domesticated yeast *Saccharomyces cerevisiae* experienced significantly increased fitness after only tens of generations, or a few weeks, in a novel chemostat environment (Goddard, Godfray, & Burt, 2005). Similarly, rapid microbial adaptation is well documented in other human-influenced systems, including antibiotic-treated patients in hospitals, crop systems with microbial pathogens, domesticated fermentations, and the human microbiome (Beaume et al., 2017; Davies & Davies, 2010; McDonald & Stukenbrock, 2016; Verstrepen, Derdelinckx, Verachtert, & Delvaux, 2003). Observations of rapid adaptation in human-influenced microbial populations may or may not extend to wild microbial populations because humans influence the ecology of their associated microbial populations (Gibbons & Rinker, 2015). For example, humans generally interact with rapidly multiplying microbial populations under low dispersal limitation (Beggs, Knibbs, Johnson, & Morawska, 2015; Kawecki et al., 2012). Studies of microbial diversity and fitness *in realistic natural environments* are needed to understand microbial responses to their changing native habitats.

Populations of *Saccharomyces paradoxus*, the wild sister species of *S. cerevisiae*, are an ideal model system for investigating adaptive evolution in nature. Both *S. paradoxus* and *S. cerevisiae* evolve quickly in laboratory evolution experiments (Goddard & Bradford, 2003; Goddard et al., 2005; Ratcliff, Denison, Borrello, & Travisano, 2012; Selmecki et al., 2015; Wloch-Salamon, Fisher, & Regenberg, 2017). Unlike *S. cerevisiae, S. paradoxus* has never been domesticated; it lives in forest environments, primarily in the northern hemisphere (Boynton & Greig, 2014; Robinson, Pinharanda, & Bensasson, 2016). It associates with oak trees, and can be found on bark, exudates, leaf litter, and soil near oaks (Boynton, Kowallik, Landermann, & Stukenbrock, 2019; Kowallik & Greig, 2016; Kowallik, Miller, & Greig, 2015; Naumov, Naumova, & Sniegowski, 1998; Sniegowski, Dombrowski, & Fingerman, 2002). While the range of *S. paradoxus* is limited by maximum summer temperature, population sizes can be stable year-round through seasonal changes in temperature, moisture, nutrient availability, and the activity of other organisms (Kowallik & Greig, 2016; Robinson et al., 2016; Voříšková, Brabcová, Cajthaml, & Baldrian, 2014). In our study forest in northern Germany, the number of *S. paradoxus* cells per gram of leaf litter does not vary from season to season, suggesting that *S. paradoxus* dormancy does not follow seasonal cycles, and that cells are instead as active in winter as they are in summer (Kowallik & Greig, 2016).

Seasonal abiotic changes, and especially temperature changes, are a plausible selective pressure on forest *S. paradoxus*. Temperature is an important selective pressure across *Saccharomyces* species: different species have different temperature optima and environmental temperatures determine *Saccharomyces* species ranges (Robinson et al., 2016; Salvadó et al., 2011; Sampaio & Gonçalves, 2008; Sweeney, Kuehne, & Sniegowski, 2004). In laboratory experimental evolution, *S. cerevisiae* adapts to previously lethal temperatures after a few days (Huang, Lu, Chang, & Li, 2018); this rapid laboratory adaptation suggests that temperature adaptation is possible on seasonal timescales in wild *Saccharomyces* species. Rapid adaptation to seasonal changes has also previously been observed in wild animal populations, which have longer generation times than laboratory *Saccharomyces*. For example, differences in fitness among Leopard Frog morphs between winter and summer lead to seasonal differences in allele frequencies (Merrell & Rodell, 1968), and stress tolerant *Drosophila* genotypes are more common in the spring (immediately after winter) than the fall (immediately after summer) (Behrman, Watson, O’Brien, Heschel, & Schmidt, 2015).

As with abiotic environmental changes, biotic interactions can also select organisms over short time scales. For example, laboratory host-pathogen and predator-prey coevolution experiments often result in rapid coadaptation among organisms as diverse as animals, bacteria, and algae. Nematodes adapt to pathogenic bacteria over tens of generations by increasing sexual recombination even as their pathogens evolve greater infectivity (Morran, Schmidt, Gelarden, Parrish, & Lively, 2011), and green algae adapt their morphologies to rotifer predators over tens of days (Becks, Ellner, Jones, & Hairston, 2010). Yeast species also rapidly adapt to biotic selective pressures during laboratory evolution. For example, bacterial communities select for diverse *S. cerevisiae* multicellular phenotypes after three months of laboratory evolution (Quintero-Galvis et al., 2018). Bacteria also select for increased fermentation ability and temperature tolerance in *Lachancea* species (*Lachancea* is a genus of yeasts in the same family as *Saccharomyces*) after months of experimental evolution (Zhou, Ishchuk, Knecht, Compagno, & Piškur, 2019; Zhou et al., 2017). Intraspecific interactions may be equally important for adaptation, as demonstrated by the evolution of *S. cerevisiae* resistance to intraspecific toxins over days of laboratory evolution (Pieczynska, Wloch-Salamon, Korona, & de Visser, 2016).

Intraspecific interference competition, mediated by “killer toxins”, is a plausible biotic selective pressure on forest *S. paradoxus*. Killer toxins are secreted yeast toxins that kill nearby sensitive yeast cells, often of the same species, and are often coded on cytoplasmic viruses (Boynton, 2019; Magliani, Conti, Gerloni, Bertolotti, & Polonelli, 1997; Schmitt & Breinig, 2006). These toxins are well studied in *S. cerevisiae* and have been reported in *S. paradoxus* (Chang, Leu, & Chang, 2015; Pieczynska, de Visser, & Korona, 2013). However, the primary purpose of killer toxins in nature is unclear: these toxins may promote interference competition in nature, maintain viruses in host cells, mediate cell-cell communication, or have another, as yet undetermined, function (Boynton, 2019). Toxin production may also be influenced by the abiotic environment. Viruses coding for killer toxins are occasionally lost after heat treatments in the laboratory (Wickner, 1974); if heat also causes virus loss in natural environments, the killer phenotype may be more common in cooler months than in warmer months. Yeasts that produce killer toxins (“killer yeasts”) select for toxin resistance in sensitive yeasts after a few days, or tens of generations, of evolution in the laboratory (Pieczynska et al., 2016). If killer toxin-mediated interference competition is an important selective pressure in nature, *S. paradoxus* populations may be under considerable pressure to evolve resistance to these toxins. If selection by killer toxins shapes forest *S. paradoxus* populations, we would expect selection for resistance to local killer toxins evolve rapidly, perhaps over seasonal timescales.

For this study, we investigated the potential for monthly-scale adaptation to seasonal environmental changes and killer toxins among members of a German *S. paradoxus* population. We had previously determined that *S. paradoxus* isolates from this forest are genetically and phenotypically diverse (Boynton et al., 2019). For the current study, we measured the population structure and killer-related phenotypes of a collection of 192 *S. paradoxus* isolates, collected over four seasons, to understand whether seasonal environments correlate with population structure and whether co-adaptation among isolates evolves over seasons. First we determined the multilocus genotype of each *S. paradoxus* forest isolate. Then we assayed each *S. paradoxus* forest isolate for killer toxin production, and investigated whether these killer yeasts selected for resistant *S. paradoxus* strains in their local environments. We tested for selection by toxins over time, expecting resistance after a killer strain was detected (but not before); and over space, expecting resistance close to a killer strain (but not far away), if these toxins selected for resistance (Figures 1a-b, 2a-b).

**Figure 1:**
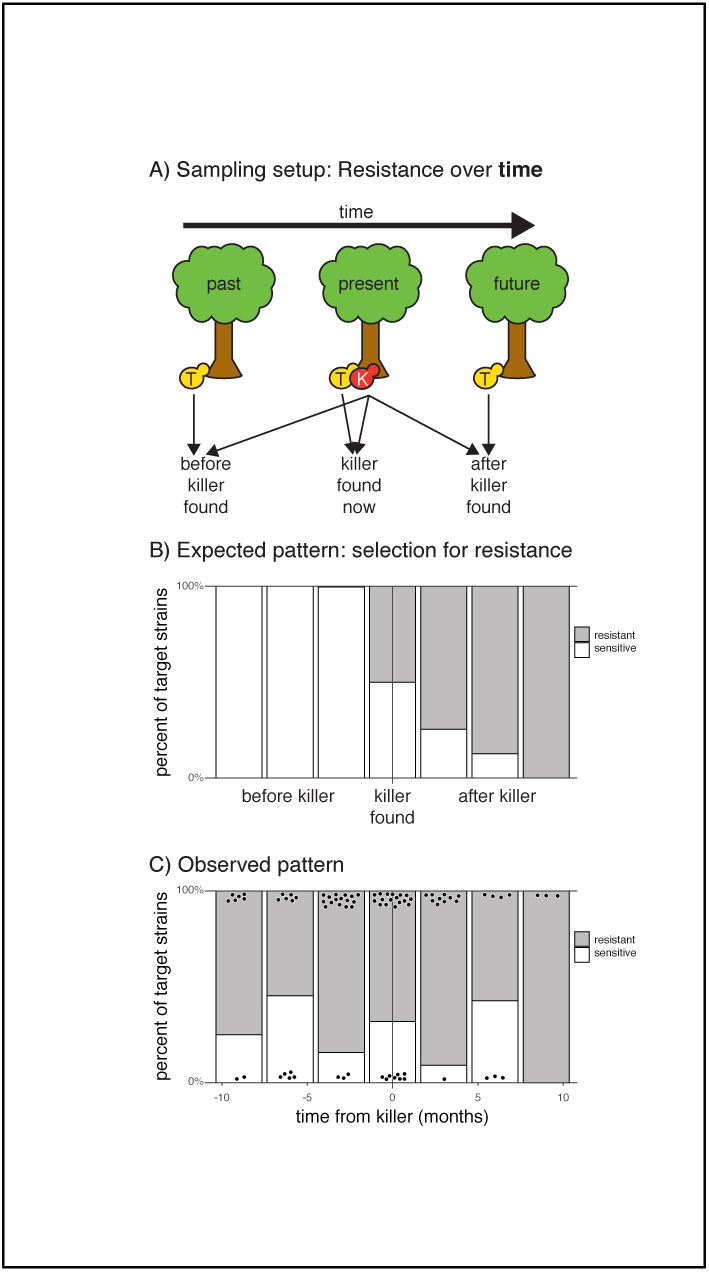
Resistance to killer toxins over time. a) Schematic of experimental design. For each toxin-producing killer strain (“K” in red), target strains (“T” in yellow) were selected from before, during, and after the killer strain was found, as available. Each target strain was tested for resistance or sensitivity to the killer strain. b) Cartoon of expected results if presence of a toxin-producing strain selects for resistance over time. The x-axis depicts the time in which each target yeast was found relative to the time in which its corresponding killer was found. The y-axis depicts the expected frequencies of resistant strains before, while, and after a killer was found c) Observed proportion of resistant strains over time. Gray bars indicate proportion of resistant strains; white bars indicate proportion of sensitive strains; and black points indicate individual tested target strains.

**Figure 2:**
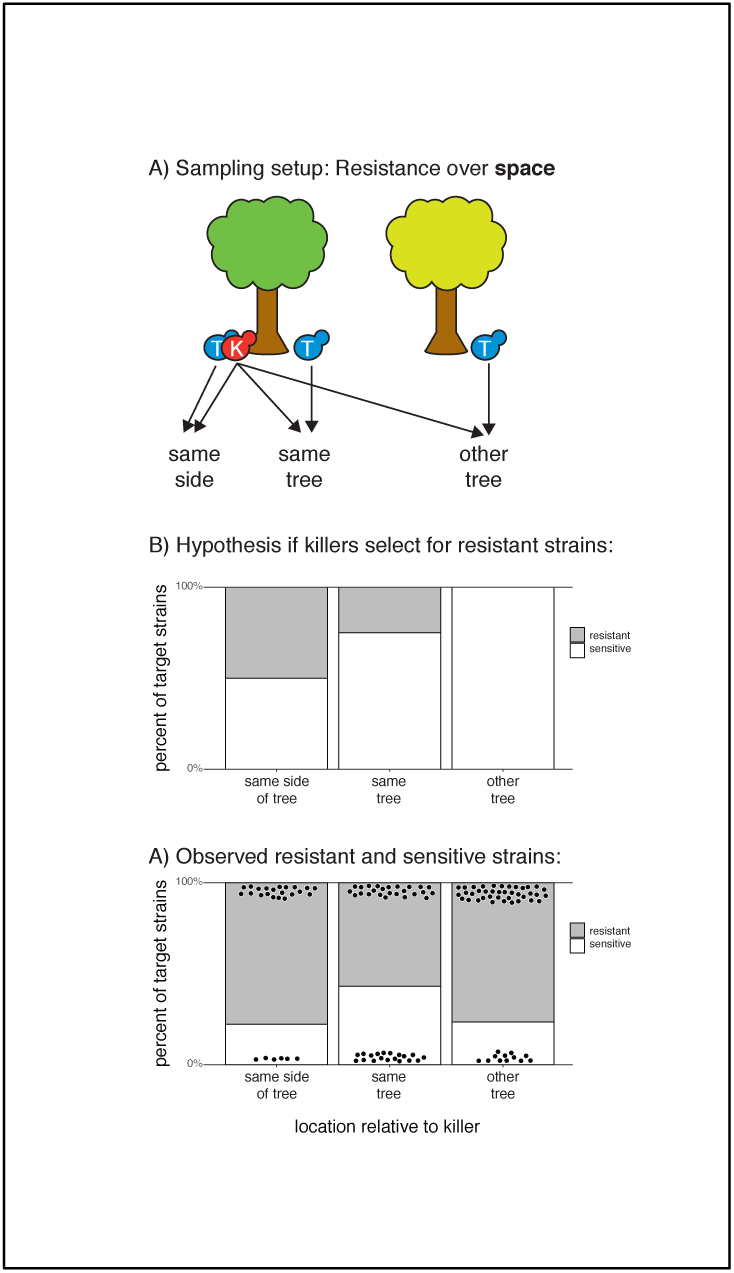
Resistance to killer toxins over space. a) Schematic of experimental design. For each toxin-producing killer strain (“K” in red), target strains (“T” in blue) were selected from the same location, another location at the same tree, and a randomly selected other sampled tree. Each target strain was tested for resistance or sensitivity to the killer strain. b) Cartoon of expected results if presence of a toxin-producing strain selects for resistance over space. The x-axis depicts the location from which each target yeast was found relative to its corresponding killer’s location. The y-axis depicts the expected frequencies of resistant strains before, while, and after a killer was found c) Observed proportion of resistant strains over space. Gray bars indicate proportion of resistant strains; white bars indicate proportion of sensitive strains; and black points indicate individual tested target strains.

## Methods

### S. paradoxus collection and identification from a forest floor environment

We collected *S. paradoxus* isolates from a northern German forest over four seasons in 2017-2018. Strains were collected on 12 June 2017 (late spring), 9 September 2017 (late summer), 8 December 2017 (late fall), and 22 March 2018 (late winter) from soil and leaf litter next to eight oak trees in the Nehmtener Forst, Nehmten, Schleswig-Holstein, Germany (approximate latitude 54.1, longitude 10.4) (Table 1). The trees were between 12 and 746 meters from one another. We collected strains using the direct plating protocol described previously (Boynton et al., 2019). Briefly, we collected samples of approximately 5 cm^3^ compressed leaf litter and soil from four locations evenly spaced around each tree base on north, south, east, and west sides of the tree. Each sample was collected within 1 m of the tree base, and repeated samplings from the same side of each tree were within approximately 50 cm of one another. We mixed each sample with 10 ml sterile water, vigorously vortexed the mixture, and spread 200 µl onto each of two solid selective plates (3 g yeast extract, 5 g peptone, 10 g sucrose, 3 g malt extract, 1 mg chloramphenicol, 80 ml ethanol, 5.2 ml 1 M HCl, and 20 g agar per liter) (Sniegowski et al., 2002). After three days, up to twelve yeast colonies with *Saccharomyces*-like morphology (round, cream-colored colonies) were randomly picked and stored in 20% glycerol at −70 to −80 °C.

**Table 1:**
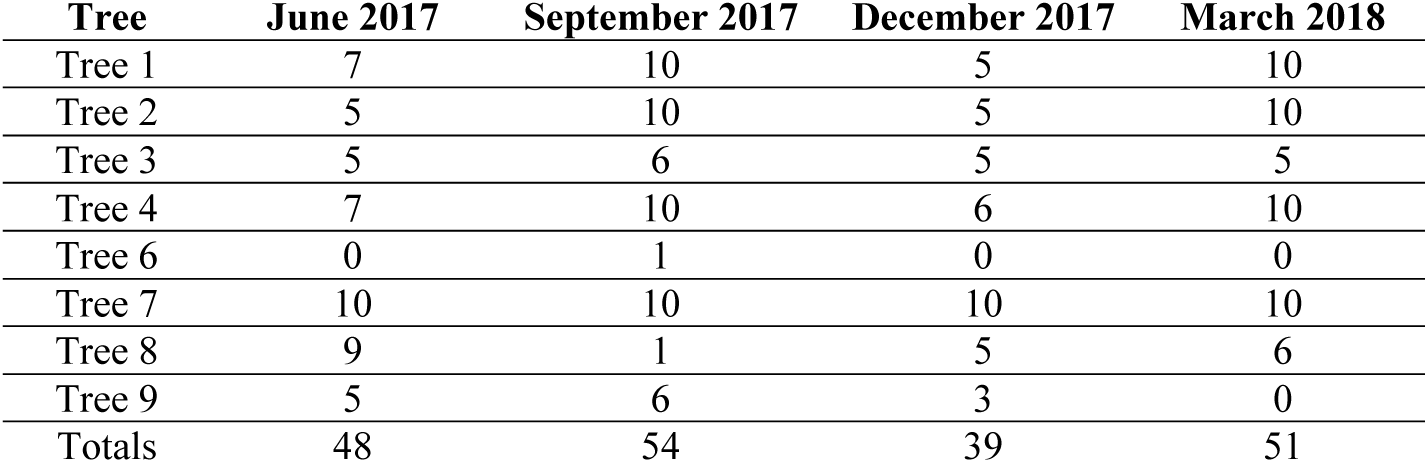
number of samples genotyped at each tree and date.

We used morphological and mating screenings to determine whether yeast colonies were *S. paradoxus* as described previously (Boynton et al., 2019). Colonies were first screened for the ability to form *Saccharomyces*-like asci (“tetrads”) on sporulation medium (20 g potassium acetate, 2.2g yeast extract, 0.5 g dextrose, 870 mg complete amino acid mixture, and 25 g agar per liter). Colonies that produced tetrads were then mated with the *S. paradoxus* tester strain NCYC 3708. Because haploid spores from different *Saccharomyces* species can mate to form zygotes (Greig, Borts, Louis, & Travisano, 2002), this mating test only identifies isolates to genus. However, the vast majority (over 99%) of *Saccharomyces* isolates from the Nehmten forest were identified as *S. paradoxus* in a previous study (Boynton et al., 2019), and we therefore assumed that all isolates that successfully mated with NCYC 3708 were *S. paradoxus*.

We randomly selected up to five *S. paradoxus* isolates from each environmental combination of collection month, source tree, and source substrate (leaf litter or soil) for further genotype and phenotype tests. Some environmental combinations produced less than five isolates, and we included all available isolates for these combinations. In total, a collection of 192 isolates were used for genotyping, killer toxin production screening, and toxin resistance screening (Table 1).

### Microsatellite genotyping

We genotyped all 192 *S. paradoxus* forest isolates plus 23 *S. paradoxus* isolates from the *Saccharomyces* Genotype Resequencing Project (SGRP) using microsatellite loci and protocols described previously (Babiker & Tautz, 2015; Boynton et al., 2019; Hardouin et al., 2015). SGRP strains were included to put local forest *S. paradoxus* into a population context, and included 22 strains from the European *S. paradoxus* population and one from the Far Eastern *S. paradoxus* population, which we used as an outgroup (Liti et al., 2009). Lengths of nine microsatellite loci on six chromosomes were determined for all strains (Table 2) (Boynton et al., 2019). We amplified loci in 5 µl multiplex PCR reactions containing four or five primer pairs each; reactions were composed of one *S. paradoxus* colony, 2.5 µl Qiagen Multiplex PCR master mix, and 0.2 µM each primer in PCR-grade water. Forward primers were labeled with FAM, HEX, or NED fluorophores at the 5’ end. PCR cycling, dilution, and denaturation were carried out using previously described protocols (Babiker & Tautz, 2015; Hardouin et al., 2015). We determined fragment lengths using an ABI 3730 DNA analyzer, Geneious 8.1.8 (https://www.geneious.com), and the Geneious microsatellite plugin 1.4.4, with Genescan ROX-550 as a size standard, as previously described (Boynton et al., 2019).

**Table 2:**
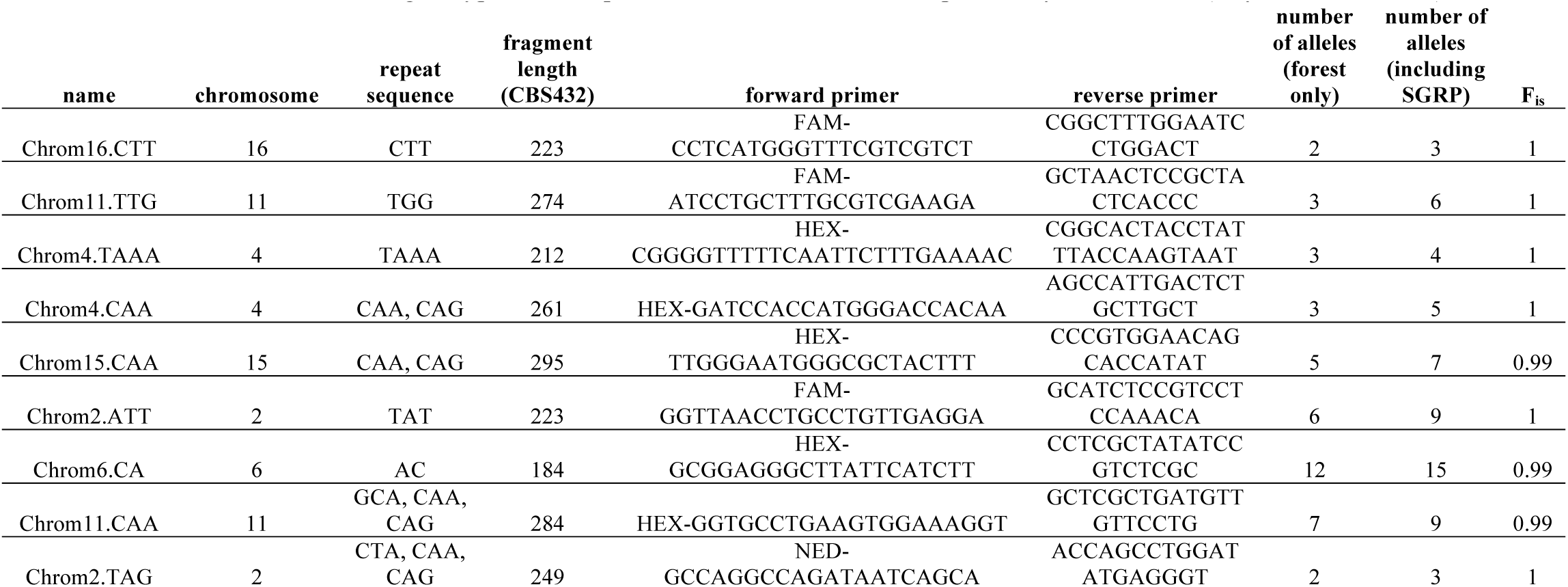
Microsatellite loci used to genotype forest *S. paradoxus* isolates. Loci were previously described in (Boynton et al., 2019).

### Screening for killing ability

Each of the 192 forest *S. paradoxus* strains was tested for the ability to kill five sensitive tester *S. cerevisiae* strains using “halo assays” (Boynton, 2019; Chang et al., 2015; Woods & Bevan, 1968). To conduct a halo assay, drops of liquid containing potential killer cells were pipetted onto a lawn of potentially sensitive cells; a zone of inhibition (a “halo”) was observed if a toxin produced by drop cells killed or inhibited lawn cells (Figure S1a). Tester *S. cerevisiae* strains included Y55, s288c, and a previously engineered sensitive strain derived from BY4741 and BY4742 (WS-29-10) (Wloch-Salamon, Gerla, Hoekstra, & de Visser, 2008). We also tested whether forest *S. paradoxus* killed two isolates of a K2 killer strain (Wickner strain 1387) that had been cured of its toxin-producing virus by incubation at 37 or 40 °C (Wickner, 1974, 1980). Curing success was confirmed experimentally by performing a halo assay using the tester *S. cerevisiae* strain WS-29-10.

To perform halo assays, each tester *S. cerevisiae* strain was first evenly spread onto Killer Test Medium plates (10 g yeast extract, 20 g peptone, 20 g glucose, 15 g agar per liter, with 0.01-0.03% methylene blue, adjusted to a pH of approximately 4.6 with citric-phosphate buffer) at a final concentration of approximately 10^4^ cells per ml of medium. After tester *S. cerevisiae* lawns dried, each forest *S. paradoxus* isolate was spotted onto the tester lawn in a 4 µl drop containing approximately 8 × 10^5^ stationary-phase *S. paradoxus* cells. As controls, we also included drops of *S. cerevisiae* killers K1, K2, and K28 on each plate. Plates were incubated at 23 °C for 4-6 days. We scored each *S. paradoxus* drop for a surrounding halo in which the tester *S. cerevisiae* strain did not grow. Often, dead *S. cerevisiae* cells at the edge of the halo appeared dark blue as a result of staining with methylene blue. Each test was conducted twice, and a forest *S. paradoxus* strain was scored as a “killer” if a halo was observed on at least one of the *S. cerevisiae* tester strains for at least one of the two tests.

### Screening for evolution of killer resistance using eclipse assays

To understand whether toxin-producing strains select for resistance to toxins in nature, we tested “target” non-killer isolates, found at the same location or time as a killer strain, for sensitivity or resistance to toxins produced by the killer strain (Figs 1a, 2a). For each killer isolate identified in the screen for killing ability, we assigned two collections of target (*i.e*., potentially sensitive or resistant) forest *S. paradoxus* isolates from the full collection of 192 *S. paradoxus* isolates: a *temporal* collection and a *spatial* collection. The *temporal* collection of target isolates was composed of all strains collected at the same location (the same side of the same tree) as the killer at all available timepoints (Figure 1a). Because killers were discovered from all four timepoints, in some cases the collection of target isolates for a particular killer strain included only isolates found before or only isolates found after the killer, but all temporal collections included target isolates from at least three timepoints. The *spatial* collection of target isolates was composed of all isolates collected from the same tree and date as the killer strain, including isolates collected from the same side of the tree and other sides of the tree, plus five randomly-chosen isolates from other trees collected on the same date as the killer (Figure 2a). For each killer strain, we assayed the sensitivity or resistance of every target strain in the killer’s temporal and spatial collection.

We assayed each target forest *S. paradoxus* isolate for resistance to its assigned forest *S. paradoxus* killer using “eclipse assays” (Figure S1b) (Kishida et al., 1996). Eclipse assays are similar to halo assays, except that the killer spot is dropped onto a small spot of the target *S. paradoxus* isolate instead of a lawn of tester *S. cerevisiae* cells. Positive eclipse assays visually resemble partial solar eclipses, with presence of an empty or blue semicircle indicating a killing reaction (Figure S1b). Eclipse assays were carried out on “eclipse media” at two different pH values (10 g yeast extract, 20 g peptone, 20 g glucose, 25 g agar, and 0.008% methylene blue per liter, adjusted to a pH of 4.1 or 3.6 with 12 ml or 16 ml 1 M citric acid per liter, respectively). Each test for resistance was carried out twice, once at each pH. Approximately 100 cells of each target isolate were spotted onto each eclipse medium plate in a volume of 5 µl. After spots had dried, each spot was overlaid with approximately 10^6^ killer forest *S. paradoxus* cells in a volume of 0.5 µl. Each plate also contained control target isolate spots which were not overlaid with a killer drop. Plates were incubated at 13 °C for six days before being scored for evidence of killer toxin inhibition. We scored target strains as sensitive to a particular killer if it showed a positive reaction (zone of blue or dead cells) on either of the two tested media (pH = 4.1 or 3.6), and resistant if no positive reaction was observed.

### Data analysis

A Neighbor-joining tree of microsatellite peak data was produced from Edwards distances using the adegenet 2.1.1 and poppr 2.8.3 packages in R 3.6.0 (Edwards, 1971; Jombart, 2008; Jombart & Ahmed, 2011; Kamvar, Brooks, & Grünwald, 2015; Kamvar, Tabima, & Grünwald, 2014; R Core Team, 2019). We tested for correlations between genetic distance among strains and difference in isolation time, and correlations between genetic and geographic distances, using Mantel tests in vegan 2.5-6 (Oksanen et al., 2019). Expected heterozygosities of the forest population and the European SGRP strains were calculated using adegenet 2.1.1. We visualized the phylogenetic tree using the ggplot2 3.2.1 and ggtree 1.16.6 libraries (Wickham, 2016; Yu, Lam, Zhu, & Guan, 2018; Yu, Smith, Zhu, Guan, & Lam, 2017). The relationship between sampling effort and the number of unique genotypes observed, including a 95% confidence interval, was visualized using the interpolation method described by (Chao et al., 2014) with 50 bootstrap samplings, using iNEXT 2.0.19 (Hsieh, Ma, & Chao, 2019).

Linkage disequilibrium was calculated using the Index of Association (I_a_), determined by comparing associations between alleles according to the following equation: I_a_ = (V_o_ / V_e_) −1 (J. M. Smith, Smith, O’Rourke, & Spratt, 1993), where V_o_ is the observed variance in number of loci at which pairs of individuals in the dataset differ and V_e_ is the expected variance in the number of loci at which pairs of individuals in the dataset differ. An I_a_ of zero indicates no association among loci, and I_a_ values significantly higher than zero indicate non-random associations among loci (*i.e*., some loci appear together in the same genome more often than expected by chance) (J. M. Smith et al., 1993). Significance was determined by shuffling alleles at each locus 999 times while preserving heterozygosity and allelic structure. We calculated I_a_ for the entire forest *S. paradoxus* dataset (no correction for clonality) and for the dataset containing one example of each unique genotype (correction for clonality) using poppr. F_is_ was calculated at each locus using the pegas 0.11 package in R (Paradis, 2010).

We modeled the target strains’ resistances to killer toxins as functions of distance in time or space between when and where target and killer strains were found (Figures 1, 2). We produced separate mixed-effects logistic regressions for the temporal and spatial datasets. The response variable for both regressions was whether or not a target *S. paradoxus* strain was resistant to its killer. The fixed predictor variables were difference in time between when the killer and target strains were found for the temporal dataset, and the categorical difference in location between where the killer and target strains were found (*i.e*., same side of the same tree, different side of the same tree, or different trees) for the spatial dataset. Random effects for both regressions were the identities of the killer and target strains because not all killers were tested against all target strains. Models were constructed and tested in R using the lme4 1.1-21 and car 3.0-3 packages (Bates, Mächler, Bolker, & Walker, 2015; Fox & Weisberg, 2019).

## Results

### Stable S. paradoxus diversity over space and time

There were 41 unique genotypes among the 192 forest *S. paradoxus* isolates (Figures 3, 4, S3). However, our sampling did not saturate *S. paradoxus* diversity, as indicated by an increasing relationship between analyzed isolates and observed unique genotypes (Figure S2), and there is further unsampled diversity in the forest. Among the sampled forest isolates, sixteen genotypes were represented by multiple individuals, and no forest *S. paradoxus* isolate had the same genotype as a reference SGRP individual. The number of alleles per locus ranged from 2 to 12 (forest isolates only) or 3 to 15 (all isolates) (Table 2). We assume all individuals sharing a multilocus genotype were clones of one another because the relationship between number of loci sampled and unique genotypes detected plateaued between eight and nine loci (Figure S4). Over half of the forest *S. paradoxus* isolates (104) were members of a single genotype, and we assume these were all clones of one another (Figure 4); this common genotype was present at all four sampling times and on seven of the eight sampled trees (Figure 3). The expected heterozygosity, a measure of genetic diversity, of the forest *S. paradoxus* isolates was 0.41, which was less than the expected heterozygosity of measured European SGRP strains (0.60). Ten of the 192 forest strains (5 %) produced a toxin that killed at least one tester *S. cerevisiae* strain. The number of tester *S. cerevisiae* strains killed ranged from one (three forest *S. paradoxus* isolates) to all five (one isolate). This incidence of forest killer isolates is an underestimate because only killer strains that killed a tester strain under the pH and temperature conditions of our screen could be detected. Systematic tests with more tester strains and a variety of pH and temperature values are likely to uncover more killer isolates.

**Figure 3:**
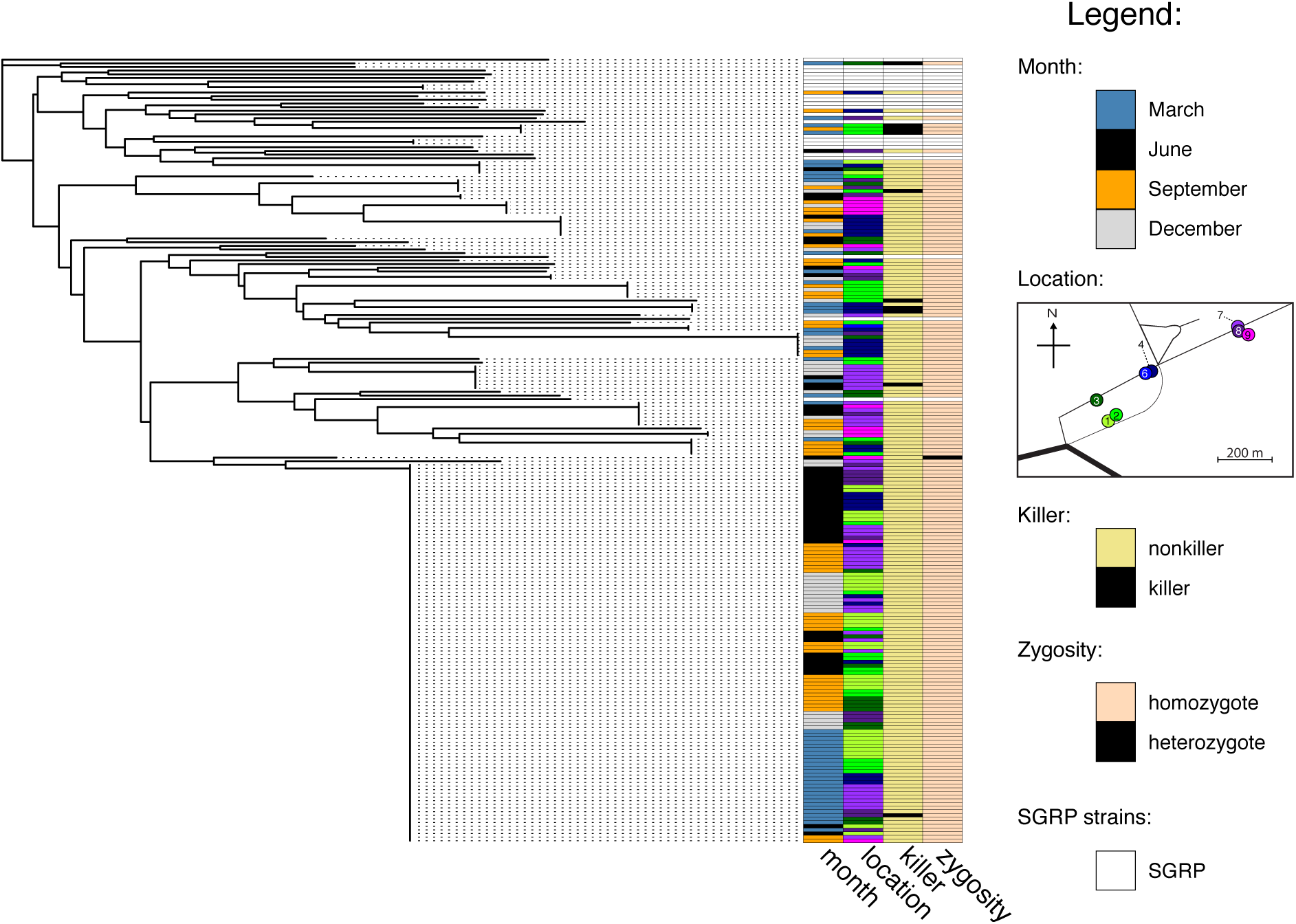
Neighbor-joining tree of all 192 forest and 23 SGRP strains genotyped, with information on the month sampled, location, ability to kill tester *S. cerevisiae*, and zygosity of each forest strain depicted with colored bars to the right of the tree. Information is not provided for SGRP strains, which are represented with white bars. See Figure S3 for a version of this figure with strain numbers at the tips.

**Figure 4:**
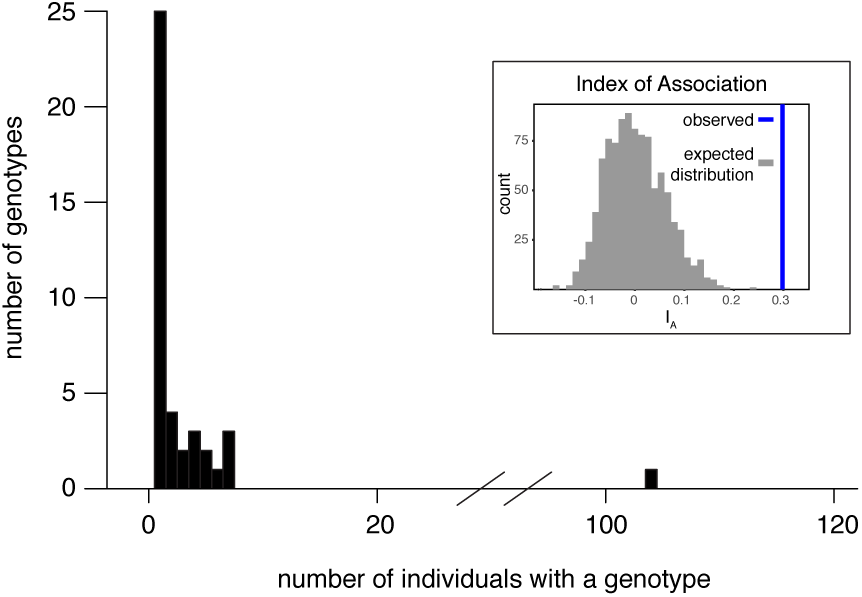
Histogram of unique observed genotypes. The x-axis depicts the number of strains in which a genotype was observed, and the y-axis depicts the number of genotypes with a given number of observed strains. The inset depicts the Index of Association (I_a_) of the clone-corrected population (*i.e*., each genotype is represented only once in the Index of Association calculation). The thick blue line represents the calculated I_a_ for the population and the gray bars represent the distribution of expected I_a_ for a population with the same allele frequencies, but randomly shuffled.

In general, the forest *S. paradoxus* population was not structured over time or space, although killer isolates were somewhat spatially restricted. There was no correlation between genetic distance and time (Mantel’s r = −0.03, p = 0.91) or space (Mantel’s r = −0.003, p = 0.57) (Figure 3). Of the 16 clones found more than once, eight were restricted to a single tree and eight were present on multiple trees. The largest clone restricted to a single tree included seven isolates (Figure 3). Of the ten detected killer *S. paradoxus* isolates, eight were found during cold months (December and March), a frequency not different from random chance (binomial test of 8 successes in 10 trials compared to .47, the proportion of total December and March isolates, p = .054) (Figure 3). Killer isolates were found at five of the eight sampled locations, with five isolates found on Tree 2 (three of these five were clones with the same microsatellite genotype). The number of isolates on Tree 2 was more than expected by random chance (binomial test of 5 successes in 10 trials compared to .16, the proportion of total strains found on Tree 2, p = .012).

In addition to its high clonality, the forest *S. paradoxus* population was characterized by high inbreeding. Only one out of the 192 isolates was heterozygous, at three of the nine loci (Figure 3, Table 2). The forest *S. paradoxus* population also had high linkage disequilibrium, as indicated by a high index of association among alleles: after correcting for clonality, the index of association among forest isolates was significantly different from zero (I_a_ = 0.30, p = .001) (Figure 4 inset). Without correcting for clonality, I_a_ was 2.87 (p = .001).

### No evidence for selection for toxin resistance

As with microsatellite genotypes, there was diversity among forest strains’ responses to killer toxins, but no evidence for selection for resistance over time or space. Resistant strains were just as common after as before the date a killer strain was found (*Χ*^2^ = 1.20, df = 1, p = .27) (Figure 1c). Similarly, resistant strains were just as common close to as far away from killer strains (*Χ*^2^ = 1.78, df = 2, p = .41) (Figure 2c). Of the 206 total killer-target combinations tested, 59 (29 %) resulted in killer inhibition of the target yeast. In agreement with previous observations of an optimal pH of approximately 4.0-4.8 for well-known *S. cerevisiae* killer toxins (McBride, Greig, & Travisano, 2008; Woods & Bevan, 1968), inhibition was more frequent on high-pH (pH = 4.1) than low-pH (pH = 3.6) media: 58 killer-target combinations showed inhibition on high-pH media, compared to 27 on low-pH media. Additionally, and anecdotally, some killer yeasts were more effective against the tested target strains than others. Of the ten killer isolates, three produced effective toxins against 75 % or more of tested target strains, while the other seven produced effective toxins against less than 9 % of tested target strains (Figures S5-S6). However, our sampling was not designed to systematically test for this effect (*i.e*., each individual killer was tested against a different set of target strains), and it is not possible to tell from these data whether the difference in apparent killing ability is an effect of killer or target strains.

## Discussion

### The forest S. paradoxus population is surprisingly robust to seasonal environmental changes

We found no evidence that seasonal-scale environmental changes select for seasonally adapted individuals from the diverse forest *S. paradoxus* population. Although we identified 41 unique *S. paradoxus* microsatellite genotypes, the genotype of an individual did not correlate with the season in which it was found. We expected potentially selected loci to be linked to our measured microsatellite loci because the *S. paradoxus* population experiences low recombination and high colonality (Figures 3, 4 inset). Similar to genotype diversity, monthly-scale selection was not associated with killer toxin-related phenotypes. *S. paradoxus* population stability is consistent with a lack of local adaptation to substrate previously observed in the same population, and with persistent North American *S. paradoxus* genomes observed in a previous study (Boynton, Stelkens, Kowallik, & Greig, 2017; Xia et al., 2017).

The forest *S. paradoxus* population’s stability contrasts with observed rapid adaptation in laboratory-selected yeast populations. *Saccharomyces* yeast populations, including *S. paradoxus* and its sister species, the laboratory model *S. cerevisiae*, have adapted over days to months to changes in temperature (Huang et al., 2018), nutrient availability (Goddard & Bradford, 2003), environmental stresses (Dettman, Sirjusingh, Kohn, & Anderson, 2007), and exposure to killer toxins (Pieczynska et al., 2016). The difference in adaptation time between forest and laboratory *S. paradoxus* populations is unlikely to be due to the number of cells in the population because forest *S. paradoxus* populations have more cells than many laboratory populations: multiplying tens to thousands of estimated *S. paradoxus* cells per gram of leaf litter (Kowallik & Greig, 2016) by metric tons of oak leaf litter per hectare per year in temperate forests (Bray & Gorham, 1964) gives 10^7^-10^9^ *S. paradoxus* cells per hectare of forest. We found little evidence for dispersal limitation over a 700 m forest transect, especially for the most common *S. paradoxus* clone (Figure 3), suggesting that there is little dispersal limitation over tens of hectares in this forest.

Nonetheless, laboratory *Saccharomyces* populations are often under selective pressures outside of the range of the environmental conditions *Saccharomyces* experiences in nature. When studying laboratory adaptation, researchers generally purposely choose selective environments that are very different from standard laboratory growth conditions (Swamy & Zhou, 2019); these selective environments are also likely to be outside of the range of environmental conditions experienced by wild yeasts. Additionally, density may be especially important for laboratory rapid adaptation. Forest *S. paradoxus* populations are considerably less dense than many adapting laboratory populations, with up to thousands of cells per gram of soil instead of as many as 10^6^ cells per ml of laboratory media (Kowallik & Greig, 2016; Zeyl, Vanderford, & Carter, 2003). Adaptation to high density can confound observations of rapid adaptation to new environmental conditions during experimental evolution.

### Population stability suggests flexibility in a seasonally changing environment

The observed *S. paradoxus* stability in the face of seasonal changes is likely the result of generalism and phenotypic plasticity evolved over thousands of years of evolutionary time. European *S. paradoxus* diverged from East Asian *S. paradoxus* tens of thousands to hundreds of thousands of years ago (Liti, Barton, & Louis, 2006), and *S. paradoxus* could have been living in northern Germany as early as the end of the last glaciation, approximately 18-19,000 years ago (Stroeven et al., 2016). *S. paradoxus* most likely adapted to seasonal fluctuations during this time, and is likely to be phenotypically plastic with regard to the biotic and abiotic variation it normally encounters over the course of months. This adaptation is further supported by high resistance to naturally-occurring killer toxins (Figures 1c, 2c), which suggests that sensitive *S. paradoxus* adapted to the presence of toxin-producing *S. paradoxus* many generations ago. Toxin-encoding viruses are in turn maintained in the population, despite widespread toxin resistance, if the viruses are not costly, or if they behave as selfish genetic elements (Boynton, 2019; Kast et al., 2015). The laboratory evidence for costs of hosting killer toxin-producing viruses is mixed, and depends on the yeast species investigated and investigation method used (Pieczynska, Korona, & De Visser, 2017; Wloch-Salamon et al., 2008). The ecological and evolutionary mechanisms maintaining these viruses in forest populations remain an open question.

High clonality and inbreeding support our observations of a stable forest *S. paradoxus* population. Sex is predicted to evolve, and outcrossing to be favored, in environments with coevolutionary selection pressures (Morran et al., 2011). The lack of observed recombination among forest *S. paradoxus* suggests that killer and sensitive *S. paradoxus* are not co-evolving in the forest. Conversely, sex and outcrossing are not favored in stable environments because recombination breaks up locally adapted gene combinations (Barton & Charlesworth, 1998). The observed levels of clonality and inbreeding in the Nehmtener forest are consistent with observations of other European and North American *S. paradoxus* populations (Johnson et al., 2004; Koufopanou, Hughes, Bell, & Burt, 2006; Tsai, Bensasson, Burt, & Koufopanou, 2008; Xia et al., 2017). Wild European *S. paradoxus* are estimated to outcross once in every 100 meiosis events, and that meiosis occurs only once in every 1000 cell divisions (Tsai et al., 2008). This lack of observed *S. paradoxus* recombination suggests that forest environments are stable from the point of view of generalist or plastic *S. paradoxus* individuals.

While *S. paradoxus* populations are robust to environmental changes on monthly time scales, our observations do not reject selection on shorter or longer time scales. For example, researchers working on weekly timescales have found that *S. paradoxus* forest population sizes spike after rain events (Anderson et al., 2018), and such events may impose selective pressures that were not detectable in our sampling scheme. In the other direction, there is strong evidence that *Saccharomyces* species have been adapting to changing environments over longer time scales. For example, *S. cerevisiae* domestication over hundreds to thousands of years has led to changes in copy number variation, ploidy, and aneuploidy during adaptation to diverse environments (Peter et al., 2018). Additionally, partially reproductively isolated *S. paradoxus* populations are locally adapted to climactic conditions in North America (Leducq et al., 2014). The temporal scales at which selection is most effective is emerging as an important question for wild microbial populations.

### The future of local S. paradoxus populations

An open question about microbial populations is how they will respond to anthropogenic climate change (Antwis et al., 2017), and our data suggest that European *S. paradoxus* populations will persist in the face of some forms of climate change. As long as temperatures remain within current seasonal ranges (*i.e*., maximum summer temperatures remain between 18 and 31 °C) (Robinson et al., 2016), we expect *S. paradoxus* to survive, even as climate variance is predicted to increase (Vasseur et al., 2014). Similarly, we expect *S. paradoxus* populations to be robust to changes in seasonal timing, such as earlier spring temperatures or later winter temperatures (Cleland, Chuine, Menzel, Mooney, & Schwartz, 2007). Once mean or maximum temperatures increase, however, *S. paradoxus* may be replaced by related warm-adapted yeasts. Specifically, *S. cerevisiae* is likely to replace *S. paradoxus* if maximum summer temperatures increase to be consistently above 31 °C because the ranges of both yeasts are limited by maximum summer temperatures (Robinson et al., 2016). This possibility demonstrates how ecological processes can be as important as evolutionary processes in determining population and community responses to climate change.

## Conclusions

The forest *S. paradoxus* population presents a paradox: it is characterized by stability, high dispersal, clonality, and inbreeding, while still including genetic and phenotypic diversity. In laboratory evolution, rapid adaptation can produce such diversity, but the evolutionary processes at play in the laboratory and forest are different. Our observations suggest that forest *S. paradoxus* do not repeatedly adapt to seasonal-scale selective pressures, and are instead plastic generalists. Future research is needed to unite the natural history of these wild microorganisms with physiological and genetic studies of plastic responses to environmental changes. *S. paradoxus* is an exciting opportunity to explore connections between forest and laboratory biology, and to understand how the biology of a model microbe influences its ecology and evolution—including its stability—in its native habitat.

## Acknowledgements

This work was supported by a Max Planck Fellowship to Eva H. Stukenbrock and grants from the National Science Center of Poland (OPUS grant 2017/25/B/NZ8/01035) and the Institute of Environmental Sciences, N18/DBS/00003, Jagiellonian University, to Dominika Wloch-Salamon. We thank Christoph Freiherr von Fürstenberg-Plessen for permission to work in the Nehmten forest, Duncan Greig for some *S. cerevisiae* strains, and Rahul Unni for insightful discussions about killer yeasts.

## Data Accessibility

Data for this project are available from the Edmond Open Access Data Repository (https://dx.doi.org/10.17617/3.3u). Deposited datasets include a list of *S. pardoxus* isolates with sampling data, microsatellite data for each isolate, the ability of each isolate to inhibit each tester *S. cerevisiae*, and the ability of killer *S. paradoxus* to inhibit tester *S. paradoxus* in temporal and spatial assays for killer resistance.

## Supplemental Figures

Figure S1: Examples of assays used to detect killer toxin activity and resistance to killer toxins. a) Example of a “halo assay”. The forest *S. paradoxus* strains 5088 and 5574 were dropped onto a lawn of the *S. cerevisiae* tester strain S288c. Strain 5574 induced an empty zone in which strain S288c could not grow, ringed by S288c colonies containing dead cells that dyed blue with methylene blue. Strain 5088 produced no empty zone or blue S288c colonies. b) Example of a “eclipse assay”. The top row is assay spots in which the forest killer *S. paradoxus* strain 5135 was dropped onto drops of the target forest *S. paradoxus* strains 5207 and 5206. The bottom row is control spots of 5207 and 5206. Strain 5135 induces a semicircle of small colonies that stain blue with methylene blue in strain 5207 but not 5206.

Figure S2: The relationship between sampling effort (number of individuals genotyped) and the number of unique genotypes observed. The thick line represents average genotypes observed as a function of isolates sampled and shaded areas represent 95% standard errors. The curve was calculated using the iNEXT package in R and the interpolation method described by (Chao et al. 2014), with the shaded area representing a 95% confidence error calculated with 50 replications.

Figure S3: Neighbor-joining tree of all 192 forest and 23 SGRP strains genotyped, with information on the month sampled, location, ability to kill tester *S. cerevisiae*, and zygosity of each forest strain depicted with colored bars to the right of the tree. Information is not provided for SGRP strains, which are represented with white bars. Strain numbers are at branch tips.

Figure S4: The relationship between number of microsatellite loci sampled and the total number of unique genotypes observed. Box plots indicate the medians, quartiles, and ranges of observed genotypes after randomly sampling 100 times. The dotted red line is the total number of genotypes (41) observed when measuring all nine loci.

Figure S5: Resistance to killer toxins over time. Data are the same as for Figure 1c, but points are color-coded by the identity of the killer strain tested and sorted by whether resistance or sensitivity was observed. Each point represents a test of a single pair of killer and target strains.

Figure S6: Resistance to killer toxins over space. Data are the same as for Figure 2c, but points are color-coded by the identity of the killer strain tested and sorted by whether resistance or sensitivity was observed. Each point represents a test of a single pair of killer and target strains.

## References

Anderson, J. B., Kasimer, D., Xia, W., Schröder, N. C. H., Cichowicz, P., Lioniello, S., … Kohn, L. M. (2018). Persistence of resident and transplanted genotypes of the undomesticated yeast *Saccharomyces paradoxus* in forest soil. MSphere, 3(3), e00211–18. doi: 10.1128/mSphere.00211-18

Andrews, J. E., & Brasier, A. T. (2005). Seasonal records of climatic change in annually laminated tufas: Short review and future prospects. Journal of Quaternary Science, 20(5), 411–421. doi: 10.1002/jqs.942

Antwis, R. E., Griffiths, S. M., Harrison, X. A., Aranega-Bou, P., Arce, A., Bettridge, A. S., … Sutherland, W. J. (2017). Fifty important research questions in microbial ecology. FEMS Microbiology Ecology, 93(5), fix044. doi: 10.1093/femsec/fix044

Babiker, H., & Tautz, D. (2015). Molecular and phenotypic distinction of the very recently evolved insular subspecies *Mus musculus helgolandicus* ZIMMERMANN, 1953. BMC Evolutionary Biology, 15(1), 160. doi: 10.1186/s12862-015-0439-5

Barton, N. H., & Charlesworth, B. (1998). Why sex and recombination? Science, 281(5385), 1986. doi: 10.1126/science.281.5385.1986

Bates, D., Mächler, M., Bolker, B., & Walker, S. (2015). Fitting linear mixed-effects models using lme4. Journal of Statistical Software, 1(1). doi: 10.18637/jss.v067.i01

Beaume, M., Köhler, T., Greub, G., Manuel, O., Aubert, J.-D., Baerlocher, L., … The Swiss Transplant Cohort Study. (2017). Rapid adaptation drives invasion of airway donor microbiota by *Pseudomonas* after lung transplantation. Scientific Reports, 7(1), 40309. doi: 10.1038/srep40309

Becks, L., Ellner, S. P., Jones, L. E., & Hairston, N. G. (2010). Reduction of adaptive genetic diversity radically alters eco-evolutionary community dynamics. Ecology Letters, 13(8), 989–997. doi: 10.1111/j.1461-0248.2010.01490.x

Beggs, C., Knibbs, L. D., Johnson, G. R., & Morawska, L. (2015). Environmental contamination and hospital-acquired infection: Factors that are easily overlooked. Indoor Air, 25(5), 462–474. doi: 10.1111/ina.12170

Behrman, E. L., Watson, S. S., O’Brien, K. R., Heschel, M. S., & Schmidt, P. S. (2015). Seasonal variation in life history traits in two *Drosophila* species. Journal of Evolutionary Biology, 28(9), 1691–1704. doi: 10.1111/jeb.12690

Boynton, P. J. (2019). The ecology of killer yeasts: Interference competition in natural habitats. Yeast, 36(8), 473–485. doi: 10.1002/yea.3398

Boynton, P. J., & Greig, D. (2014). The ecology and evolution of non-domesticated *Saccharomyces* species. Yeast, 31(12), 449–462. doi: 10.1002/yea.3040

Boynton, P. J., Kowallik, V., Landermann, D., & Stukenbrock, E. H. (2019). Quantifying the efficiency and biases of forest *Saccharomyces* sampling strategies. Yeast, 36(11), 657–668. doi: 10.1002/yea.3435

Boynton, P. J., Stelkens, R., Kowallik, V., & Greig, D. (2017). Measuring microbial fitness in a field reciprocal transplant experiment. Molecular Ecology Resources, 17(3), 370–380. doi: 10.1111/1755-0998.12562

Bray, J. R., & Gorham, E. (1964). Litter Production in Forests of the World. In J. B. Cragg (Ed.), Advances in Ecological Research (Vol. 2, pp. 101–157). doi: 10.1016/S0065-2504(08)60331-1

Chang, S.-L., Leu, J.-Y., & Chang, T.-H. (2015). A population study of killer viruses reveals different evolutionary histories of two closely related *Saccharomyces sensu stricto* yeasts. Molecular Ecology, 24(16), 4312–4322. doi: 10.1111/mec.13310

Chao, A., Gotelli, N. J., Hsieh, T. C., Sander, E. L., Ma, K. H., Colwell, R. K., & Ellison, A. M. (2014). Rarefaction and extrapolation with Hill numbers: A framework for sampling and estimation in species diversity studies. Ecological Monographs, 84(1), 45–67. doi: 10.1890/13-0133.1

Cleland, E. E., Chuine, I., Menzel, A., Mooney, H. A., & Schwartz, M. D. (2007). Shifting plant phenology in response to global change. Trends in Ecology & Evolution, 22(7), 357–365. doi: 10.1016/j.tree.2007.04.003

Davies, J., & Davies, D. (2010). Origins and evolution of antibiotic resistance. Microbiology and Molecular Biology Reviews, 74(3), 417. doi: 10.1128/MMBR.00016-10

Dettman, J. R., Sirjusingh, C., Kohn, L. M., & Anderson, J. B. (2007). Incipient speciation by divergent adaptation and antagonistic epistasis in yeast. Nature, 447(7144), 585–588. doi: 10.1038/nature05856

Edwards, A. W. F. (1971). Distance between populations on the basis of gene frequencies. Biometrics, 27, 873–881.

Fox, J., & Weisberg, S. (2019). An R Companion to Applied Regression (Third). Retrieved from https://socialsciences.mcmaster.ca/jfox/Books/Companion/

Gibbons, J. G., & Rinker, D. C. (2015). The genomics of microbial domestication in the fermented food environment. Genomes and Evolution, 35, 1–8. doi: 10.1016/j.gde.2015.07.003

Goddard, M. R., & Bradford, M. A. (2003). The adaptive response of a natural microbial population to carbon- and nitrogen-limitation. Ecology Letters, 6(7), 594–598. doi: 10.1046/j.1461-0248.2003.00478.x

Goddard, M. R., Godfray, H. C. J., & Burt, A. (2005). Sex increases the efficacy of natural selection in experimental yeast populations. Nature, 434(7033), 636–640. doi: 10.1038/nature03405

Gómez, P., & Buckling, A. (2011). Bacteria-phage antagonistic coevolution in soil. Science, 332(6025), 106. doi: 10.1126/science.1198767

Greig, D., Borts, R. H., Louis, E. J., & Travisano, m. (2002). Epistasis and hybrid sterility in *Saccharomyces*. Proceedings of the Royal Society of London. Series B: Biological Sciences, 269(1496), 1167–1171. doi: 10.1098/rspb.2002.1989

Hardouin, E. A., Orth, A., Teschke, M., Darvish, J., Tautz, D., & Bonhomme, F. (2015). Eurasian house mouse (*Mus musculus* L.) differentiation at microsatellite loci identifies the Iranian plateau as a phylogeographic hotspot. BMC Evolutionary Biology, 15(1), 26. doi: 10.1186/s12862-015-0306-4

Hsieh, T. C., Ma, K. H., & Chao, A. (2019). iNEXT: iNterpolation and EXTrapolation for species diversity (Version 2.0.19). Retrieved from http://chao.stat.nthu.edu.tw/blog/software-download/

Huang, C.-J., Lu, M.-Y., Chang, Y.-W., & Li, W.-H. (2018). Experimental evolution of yeast for high-temperature tolerance. Molecular Biology and Evolution, 35(8), 1823–1839. doi: 10.1093/molbev/msy077

Johnson, L. J., Koufopanou, V., Goddard, M. R., Hetherington, R., Schäfer, S. M., & Burt, A. (2004). Population genetics of the wild yeast *Saccharomyces paradoxus*. Genetics, 166(1), 43. doi: 10.1534/genetics.166.1.43

Jombart, T. (2008). adegenet: A R package for the multivariate analysis of genetic markers. Bioinformatics, 24(11), 1403–1405. doi: 10.1093/bioinformatics/btn129

Jombart, T., & Ahmed, I. (2011). adegenet 1.3-1: New tools for the analysis of genome-wide SNP data. Bioinformatics, 27(21), 3070–3071. doi: 10.1093/bioinformatics/btr521

Kamvar, Z. N., Brooks, J. C., & Grünwald, N. J. (2015). Novel R tools for analysis of genome-wide population genetic data with emphasis on clonality. Frontiers in Genetics, 6, 208–208. doi: 10.3389/fgene.2015.00208

Kamvar, Z. N., Tabima, J. F., & Grünwald, N. J. (2014). Poppr: An R package for genetic analysis of populations with clonal, partially clonal, and/or sexual reproduction. PeerJ, 2, e281–e281. doi: 10.7717/peerj.281

Kast, A., Voges, R., Schroth, M., Schaffrath, R., Klassen, R., & Meinhardt, F. (2015). Autoselection of cytoplasmic yeast virus like elements encoding toxin/antitoxin systems involves a nuclear barrier for immunity gene expression. PLoS Genetics, 11(5), e1005005–e1005005. doi: 10.1371/journal.pgen.1005005

Kawecki, T. J., Lenski, R. E., Ebert, D., Hollis, B., Olivieri, I., & Whitlock, M. C. (2012). Experimental evolution. Trends in Ecology & Evolution, 27(10), 547–560. doi: 10.1016/j.tree.2012.06.001

Kidwell, S. M. (2015). Biology in the Anthropocene: Challenges and insights from young fossil records. Proceedings of the National Academy of Sciences, 112(16), 4922. doi: 10.1073/pnas.1403660112

Kishida, M., Tokunaga, M., Katayose, Y., Yajima, H. Y., Kawamura-Watabe, A., & Hishinuma, F. (1996). Isolation and genetic characterization of pGKL killer-insensitive mutants (iki) from *Saccharomyces cerevisiae*. Bioscience, Biotechnology, and Biochemistry, 60(5), 798–801. doi: 10.1271/bbb.60.798

Koufopanou, V., Hughes, J., Bell, G., & Burt, A. (2006). The spatial scale of genetic differentiation in a model organism: The wild yeast *Saccharomyces paradoxus*. Philosophical Transactions of the Royal Society B: Biological Sciences, 361(1475), 1941–1946. doi: 10.1098/rstb.2006.1922

Kowallik, V., & Greig, D. (2016). A systematic forest survey showing an association of *Saccharomyces paradoxus* with oak leaf litter. Environmental Microbiology Reports, 8(5), 833–841. doi: 10.1111/1758-2229.12446

Kowallik, V., Miller, E., & Greig, D. (2015). The interaction of *Saccharomyces paradoxus* with its natural competitors on oak bark. Molecular Ecology, 24(7), 1596–1610. doi: 10.1111/mec.13120

Leducq, J.-B., Charron, G., Samani, P., Dubé, A. K., Sylvester, K., James, B., … Landry, C. R. (2014). Local climatic adaptation in a widespread microorganism. Proceedings of the Royal Society B: Biological Sciences, 281(1777), 20132472. doi: 10.1098/rspb.2013.2472

Liti, G., Barton, D. B. H., & Louis, E. J. (2006). Sequence diversity, reproductive isolation and species concepts in *Saccharomyces*. Genetics, 174(2), 839. doi: 10.1534/genetics.106.062166

Liti, G., Carter, D. M., Moses, A. M., Warringer, J., Parts, L., James, S. A., … Louis, E. J. (2009). Population genomics of domestic and wild yeasts. Nature, 458(7236), 337–341. doi: 10.1038/nature07743

Luria, S. E., & Delbrück, M. (1943). Mutations of bacteria from virus sensitivity to virus resistance. Genetics, 28(6), 491.

Magliani, W., Conti, S., Gerloni, M., Bertolotti, D., & Polonelli, L. (1997). Yeast killer systems. Clinical Microbiology Reviews, 10(3), 369. doi: 10.1128/CMR.10.3.369

McBride, R., Greig, D., & Travisano, M. (2008). Fungal viral mutualism moderated by ploidy. Evolution, 62(9), 2372–2380. doi: 10.1111/j.1558-5646.2008.00443.x

McDonald, B. A., & Stukenbrock, E. H. (2016). Rapid emergence of pathogens in agro-ecosystems: Global threats to agricultural sustainability and food security. Philosophical Transactions of the Royal Society B: Biological Sciences, 371(1709), 20160026. doi: 10.1098/rstb.2016.0026

Merrell, D. J., & Rodell, C. F. (1968). Seasonal selection in the leopard frog, *Rana pipiens*. Evolution; International Journal of Organic Evolution, 22(2), 284–288. doi: 10.1111/j.1558-5646.1968.tb05896.x

Meyers, L. A., & Bull, J. J. (2002). Fighting change with change: Adaptive variation in an uncertain world. Trends in Ecology & Evolution, 17(12), 551–557. doi: 10.1016/S0169-5347(02)02633-2

Morran, L. T., Schmidt, O. G., Gelarden, I. A., Parrish, R. C., & Lively, C. M. (2011). Running with the red queen: Host-parasite coevolution selects for biparental sex. Science, 333(6039), 216. doi: 10.1126/science.1206360

Nadeau, C. P., & Urban, M. C. (2019). Eco-evolution on the edge during climate change. Ecography, 42(7), 1280–1297. doi: 10.1111/ecog.04404

Naumov, G. I., Naumova, E. S., & Sniegowski, P. D. (1998). *Saccharomyces paradoxus* and *Saccharomyces cerevisiae* are associated with exudates of North American oaks. Canadian Journal of Microbiology, 44(11), 1045–1050.

Oksanen, J., Blanchet, F. G., Friendly, M., Kindt, R., Legendre, P., McGlinn, D., … Wagner, H. (2019). vegan: Community Ecology Package (Version 2.5-6). Retrieved from https://CRAN.R-project.org/package=vegan

Paradis, E. (2010). pegas: An R package for population genetics with an integrated–modular approach. Bioinformatics, 26(3), 419–420. doi: 10.1093/bioinformatics/btp696

Peter, J., De Chiara, M., Friedrich, A., Yue, J.-X., Pflieger, D., Bergström, A., … Schacherer, J. (2018). Genome evolution across 1,011 *Saccharomyces cerevisiae* isolates. Nature, 556(7701), 339–344. doi: 10.1038/s41586-018-0030-5

Pieczynska, M. D., de Visser, J. A. G. M., & Korona, R. (2013). Incidence of symbiotic dsRNA ‘killer’ viruses in wild and domesticated yeast. FEMS Yeast Research, 13(8), 856–859. doi: 10.1111/1567-1364.12086

Pieczynska, M. D., Korona, R., & De Visser, J. A. G. M. (2017). Experimental tests of host–virus coevolution in natural killer yeast strains. Journal of Evolutionary Biology, 30(4), 773–781. doi: 10.1111/jeb.13044

Pieczynska, M. D., Wloch-Salamon, D., Korona, R., & de Visser, J. A. G. M. (2016). Rapid multiple-level coevolution in experimental populations of yeast killer and nonkiller strains. Evolution, 70(6), 1342–1353. doi: 10.1111/evo.12945

Quintero-Galvis, J. F., Paleo-López, R., Solano-Iguaran, J. J., Poupin, M. J., Ledger, T., Gaitan-Espitia, J. D., … Nespolo, R. F. (2018). Exploring the evolution of multicellularity in *Saccharomyces cerevisiae* under bacteria environment: An experimental phylogenetics approach. Ecology and Evolution, 8(9), 4619–4630. doi: 10.1002/ece3.3979

R Core Team. (2019). R: A Language and Environment for Statistical Computing (Version 3.6.0). Retrieved from https://www.R-project.org/

Rafaluk-Mohr, C., Ashby, B., Dahan, D. A., & King, K. C. (2018). Mutual fitness benefits arise during coevolution in a nematode-defensive microbe model. Evolution Letters, 2(3), 246–256. doi: 10.1002/evl3.58

Rainey, P. B., & Travisano, M. (1998). Adaptive radiation in a heterogeneous environment. Nature, 394(6688), 69–72. doi: 10.1038/27900

Ratcliff, W. C., Denison, R. F., Borrello, M., & Travisano, M. (2012). Experimental evolution of multicellularity. Proceedings of the National Academy of Sciences, 109(5), 1595. doi: 10.1073/pnas.1115323109

Robinson, H. A., Pinharanda, A., & Bensasson, D. (2016). Summer temperature can predict the distribution of wild yeast populations. Ecology and Evolution, 6(4), 1236–1250. doi: 10.1002/ece3.1919

Rowe, K. C., Rowe, K. M. C., Tingley, M. W., Koo, M. S., Patton, J. L., Conroy, C. J., … Moritz, C. (2015). Spatially heterogeneous impact of climate change on small mammals of montane California. Proceedings of the Royal Society B: Biological Sciences, 282(1799), 20141857. doi: 10.1098/rspb.2014.1857

Salvadó, Z., Arroyo-López, F. N., Guillamón, J. M., Salazar, G., Querol, A., & Barrio, E. (2011). Temperature adaptation markedly determines evolution within the genus *Saccharomyces*. Applied and Environmental Microbiology, 77(7), 2292. doi: 10.1128/AEM.01861-10

Sampaio, J. P., & Gonçalves, P. (2008). Natural populations of *Saccharomyces kudriavzevii* in Portugal are associated with oak bark and are sympatric with *S. cerevisiae* and *S. paradoxus*. Applied and Environmental Microbiology, 74(7), 2144. doi: 10.1128/AEM.02396-07

Schmitt, M. J., & Breinig, F. (2006). Yeast viral killer toxins: Lethality and self-protection. Nature Reviews Microbiology, 4(3), 212–221. doi: 10.1038/nrmicro1347

Selmecki, A. M., Maruvka, Y. E., Richmond, P. A., Guillet, M., Shoresh, N., Sorenson, A. L., … Pellman, D. (2015). Polyploidy can drive rapid adaptation in yeast. Nature, 519(7543), 349–352. doi: 10.1038/nature14187

Smith, A. B., Beever, E. A., Kessler, A. E., Johnston, A. N., Ray, C., Epps, C. W., … Yandow, L. (2019). Alternatives to genetic affinity as a context for within-species response to climate. Nature Climate Change, 9(10), 787–794. doi: 10.1038/s41558-019-0584-8

Smith, J. M., Smith, N. H., O’Rourke, M., & Spratt, B. G. (1993). How clonal are bacteria? Proceedings of the National Academy of Sciences of the United States of America, 90(10), 4384–4388. doi: 10.1073/pnas.90.10.4384

Sniegowski, P. D., Dombrowski, P. G., & Fingerman, E. (2002). *Saccharomyces cerevisiae* and *Saccharomyces paradoxus* coexist in a natural woodland site in North America and display different levels of reproductive isolation from European conspecifics. FEMS Yeast Research, 1(4), 299–306. doi: 10.1016/S1567-1356(01)00043-5

Stroeven, A. P., Hättestrand, C., Kleman, J., Heyman, J., Fabel, D., Fredin, O., … Jansson, K. N. (2016). Deglaciation of Fennoscandia. Special Issue: PAST Gateways (Palaeo-Arctic Spatial and Temporal Gateways), 147, 91–121. doi: 10.1016/j.quascirev.2015.09.016

Swamy, K. B. S., & Zhou, N. (2019). Experimental evolution: its principles and applications in developing stress-tolerant yeasts. Applied Microbiology and Biotechnology, 103(5), 2067–2077. doi: 10.1007/s00253-019-09616-2

Sweeney, J. Y., Kuehne, H. A., & Sniegowski, P. D. (2004). Sympatric natural *Saccharomyces cerevisiae* and *S. paradoxus* populations have different thermal growth profiles. FEMS Yeast Research, 4(4), 521–525. doi: 10.1016/S1567-1356(03)00171-5

Tsai, I. J., Bensasson, D., Burt, A., & Koufopanou, V. (2008). Population genomics of the wild yeast *Saccharomyces paradoxus*: Quantifying the life cycle. Proceedings of the National Academy of Sciences of the United States of America, 105(12), 4957–4962. doi: 10.1073/pnas.0707314105

Vasseur, D. A., DeLong, J. P., Gilbert, B., Greig, H. S., Harley, C. D. G., McCann, K. S., … O’Connor, M. I. (2014). Increased temperature variation poses a greater risk to species than climate warming. Proceedings of the Royal Society B: Biological Sciences, 281(1779), 20132612. doi: 10.1098/rspb.2013.2612

Verstrepen, K. J., Derdelinckx, G., Verachtert, H., & Delvaux, F. R. (2003). Yeast flocculation: What brewers should know. Applied Microbiology and Biotechnology, 61(3), 197–205. doi: 10.1007/s00253-002-1200-8

Voříšková, J., Brabcová, V., Cajthaml, T., & Baldrian, P. (2014). Seasonal dynamics of fungal communities in a temperate oak forest soil. New Phytologist, 201(1), 269–278. doi: 10.1111/nph.12481

Wickham, H. (2016). ggplot2: Elegant Graphics for Data Analysis. Retrieved from https://ggplot2.tidyverse.org

Wickner, R. B. (1974). “Killer character” of *Saccharomyces cerevisiae*: Curing by growth at elevated temperature. Journal of Bacteriology, 117(3), 1356.

Wickner, R. B. (1980). Plasmids controlling exclusion of the K2 killer double-stranded RNA plasmid of yeast. Cell, 21(1), 217–226. doi: 10.1016/0092-8674(80)90129-4

Wloch-Salamon, D. M., Fisher, R. M., & Regenberg, B. (2017). Division of labour in the yeast: *Saccharomyces cerevisiae*. Yeast, 34(10), 399–406. doi: 10.1002/yea.3241

Wloch-Salamon, D. M., Gerla, D., Hoekstra, R. F., & de Visser, J. A. G. M. (2008). Effect of dispersal and nutrient availability on the competitive ability of toxin-producing yeast. Proceedings of the Royal Society B: Biological Sciences, 275(1634), 535–541. doi: 10.1098/rspb.2007.1461

Woods, D. R., & Bevan, E. A. (1968). Studies on the nature of the killer factor produced by *Saccharomyces cerevisiae*. Microbiology, 51 (1), 115–126.

Xia, W., Nielly-Thibault, L., Charron, G., Landry, C. R., Kasimer, D., Anderson, J. B., & Kohn, L. M. (2017). Population genomics reveals structure at the individual, host-tree scale and persistence of genotypic variants of the undomesticated yeast *Saccharomyces paradoxus* in a natural woodland. Molecular Ecology, 26(4), 995–1007. doi: 10.1111/mec.13954

Yu, G., Lam, T. T.-Y., Zhu, H., & Guan, Y. (2018). Two methods for mapping and visualizing associated data on phylogeny using ggtree. Molecular Biology and Evolution, 35(12), 3041–3043. doi: 10.1093/molbev/msy194

Yu, G., Smith, D. K., Zhu, H., Guan, Y., & Lam, T. T.-Y. (2017). ggtree: an r package for visualization and annotation of phylogenetic trees with their covariates and other associated data. Methods in Ecology and Evolution, 8(1), 28–36. doi: 10.1111/2041-210X.12628

Zeyl, C., Vanderford, T., & Carter, M. (2003). An evolutionary advantage of haploidy in large yeast populations. Science, 299(5606), 555. doi: 10.1126/science.1078417

Zhou, N., Ishchuk, O. P., Knecht, W., Compagno, C., & Piškur, J. (2019). Improvement of thermotolerance in *Lachancea thermotolerans* using a bacterial selection pressure. Journal of Industrial Microbiology & Biotechnology, 46(2), 133–145. doi: 10.1007/s10295-018-2107-4

Zhou, N., Swamy, K. B. S., Leu, J.-Y., McDonald, M. J., Galafassi, S., Compagno, C., & Piškur, J. (2017). Coevolution with bacteria drives the evolution of aerobic fermentation in *Lachancea kluyveri*. PloS One, 12(3), e0173318. doi: 10.1371/journal.pone.0173318

