## Supplemental Figures for "Forest *Saccharomyces paradoxus* are robust to seasonal biotic and abiotic changes"

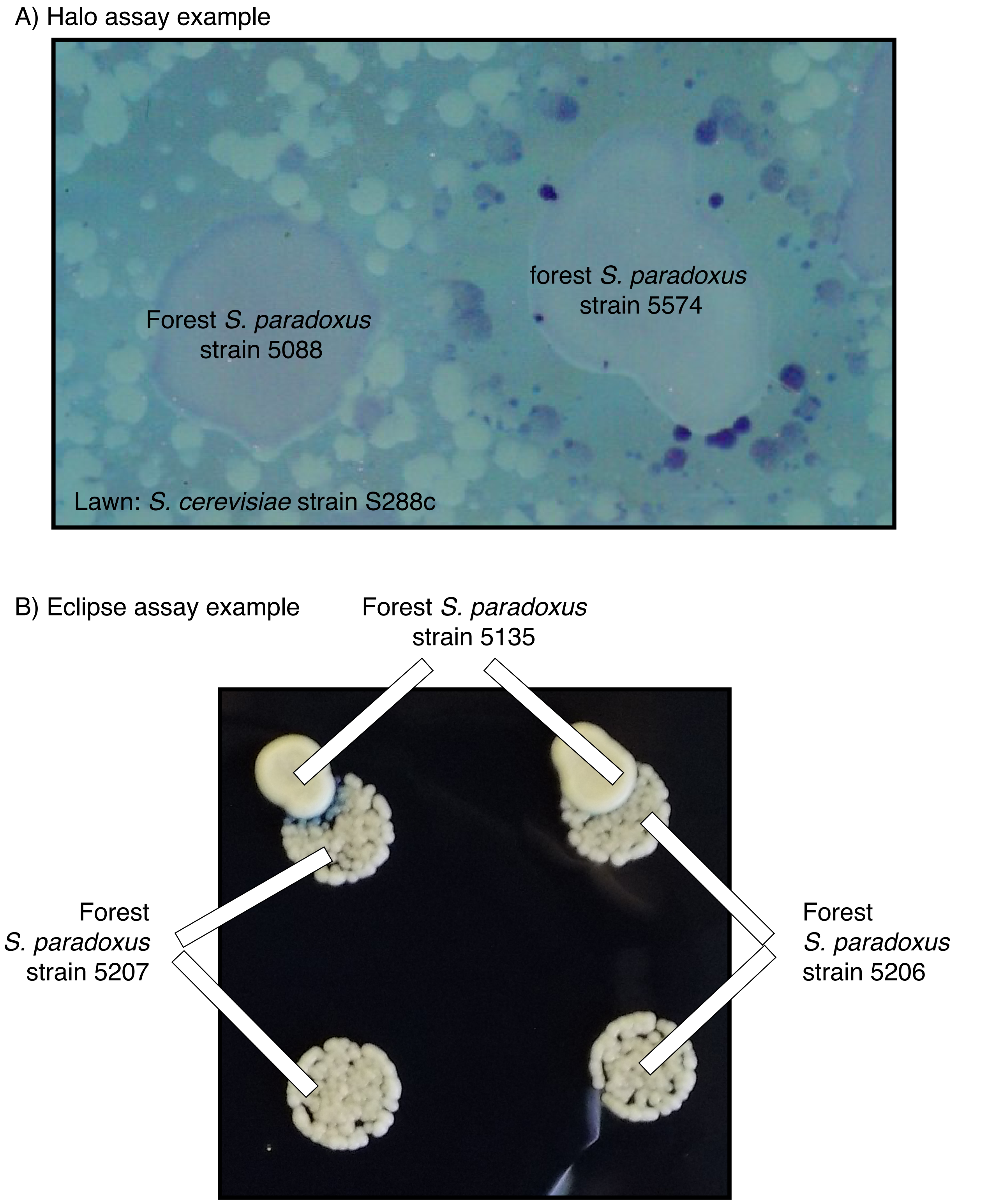


Figure S1: Examples of assays used to detect killer toxin activity and resistance to killer toxins. a) Example of a “halo assay”. The forest *S. paradoxus* strains 5088 and 5574 were dropped onto a lawn of the *S. cerevisiae* tester strain S288c. Strain 5574 induced an empty zone in which strain S288c could not grow, ringed by S288c colonies containing dead cells that dyed blue with methylene blue. Strain 5088 produced no empty zone or blue S288c colonies. b) Example of a “eclipse assay”. The top row is assay spots in which the forest killer *S. paradoxus* strain 5135 was dropped onto drops of the target forest *S. paradoxus* strains 5207 and 5206. The bottom row is control spots of 5207 and 5206. Strain 5135 induces a semicircle of small colonies that stain blue with methylene blue in strain 5207 but not 5206.

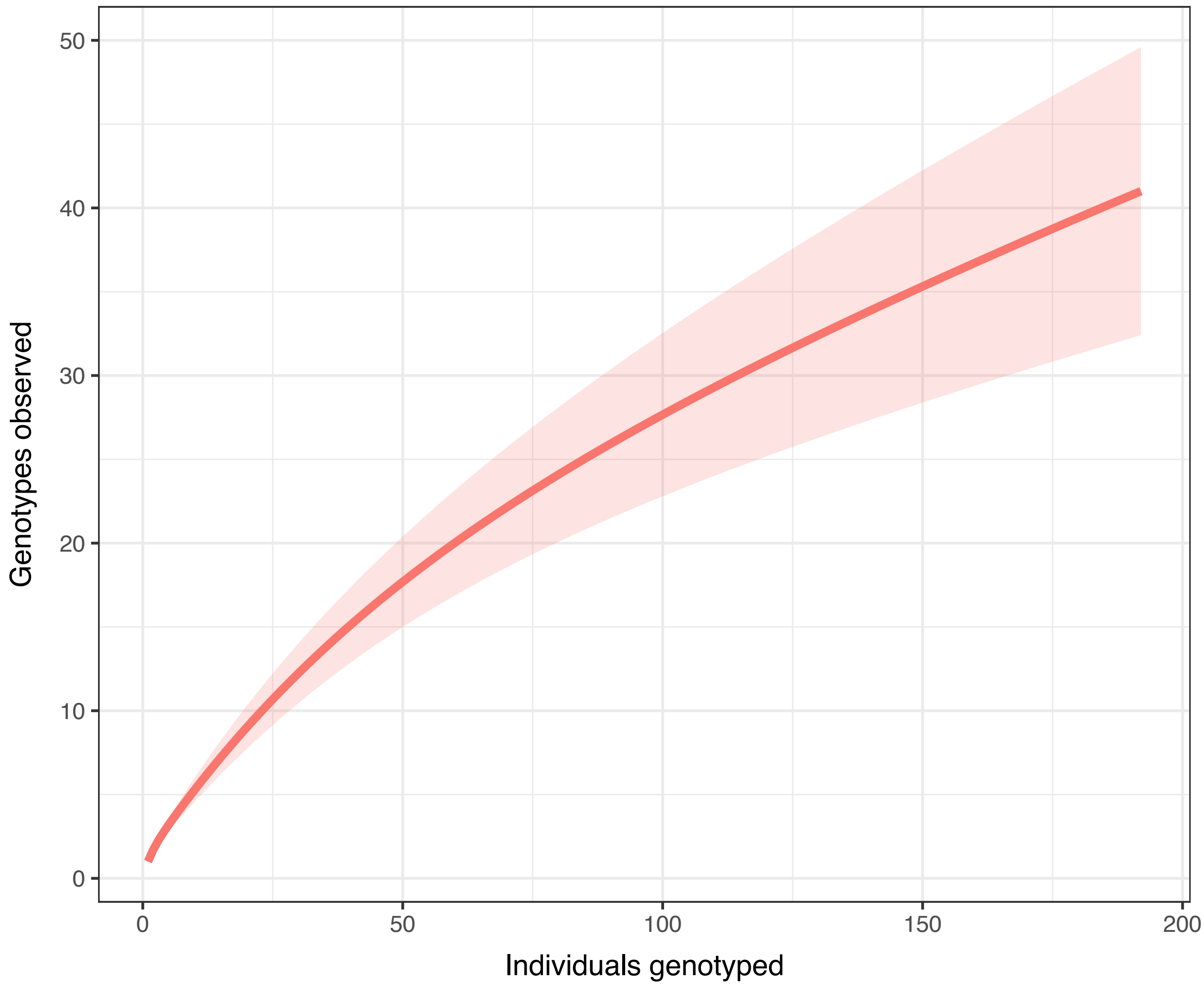


Figure S2: The relationship between sampling effort (number of individuals genotyped) and the number of unique genotypes observed. The thick line represents average genotypes observed as a function of isolates sampled and shaded areas represent 95% standard errors. The curve was calculated using the iNEXT package in R and the interpolation method described by (Chao et al. 2014), with the shaded area representing a 95% confidence error calculated with 50 replications.

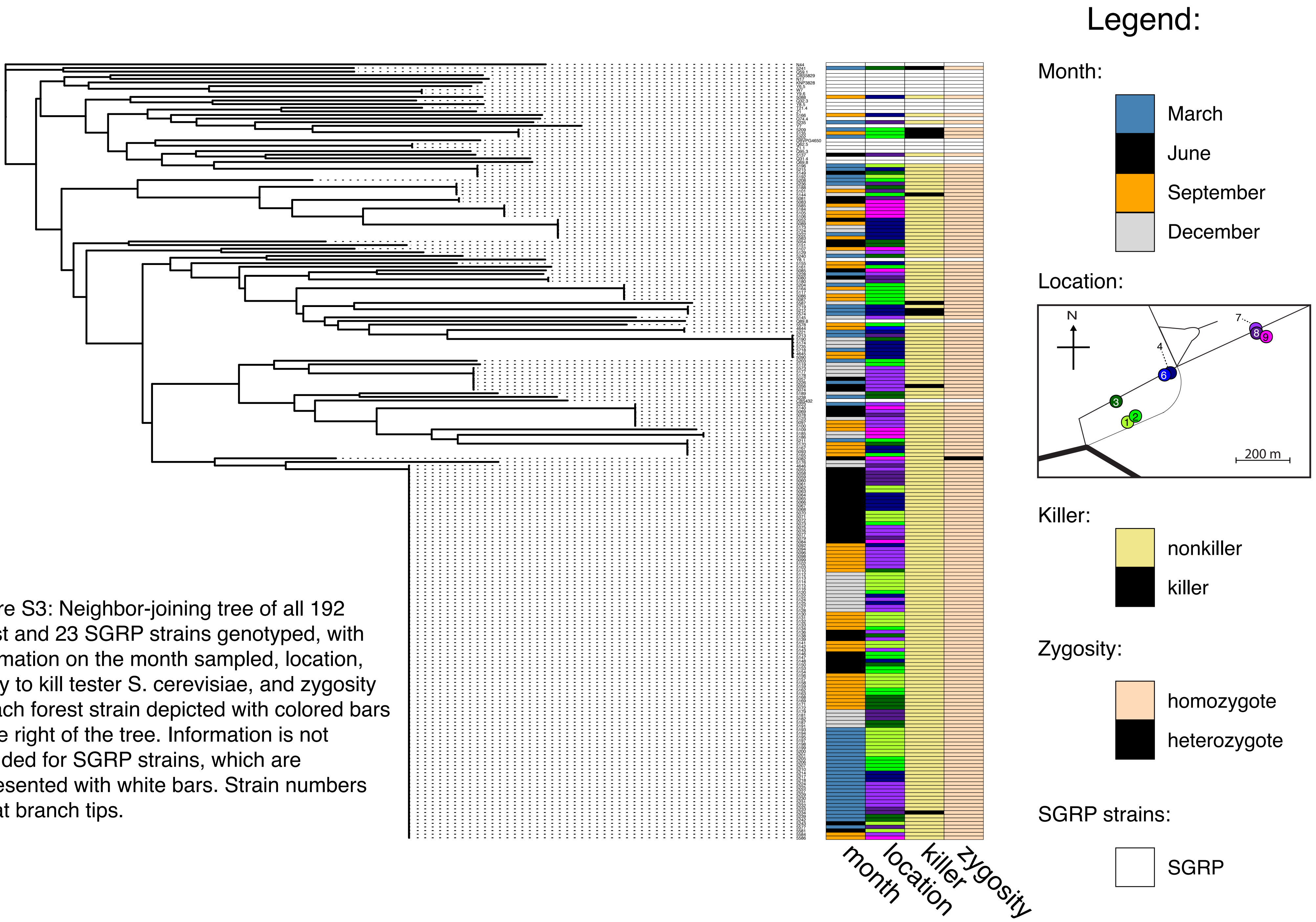


Figure S3: Neighbor-joining tree of all 192 forest and 23 SGRP strains genotyped, with information on the month sampled, location, ability to kill tester S. cerevisiae, and zygosity of each forest strain depicted with colored bars to the right of the tree. Information is not provided for SGRP strains, which are represented with white bars. Strain numbers are at branch tips.

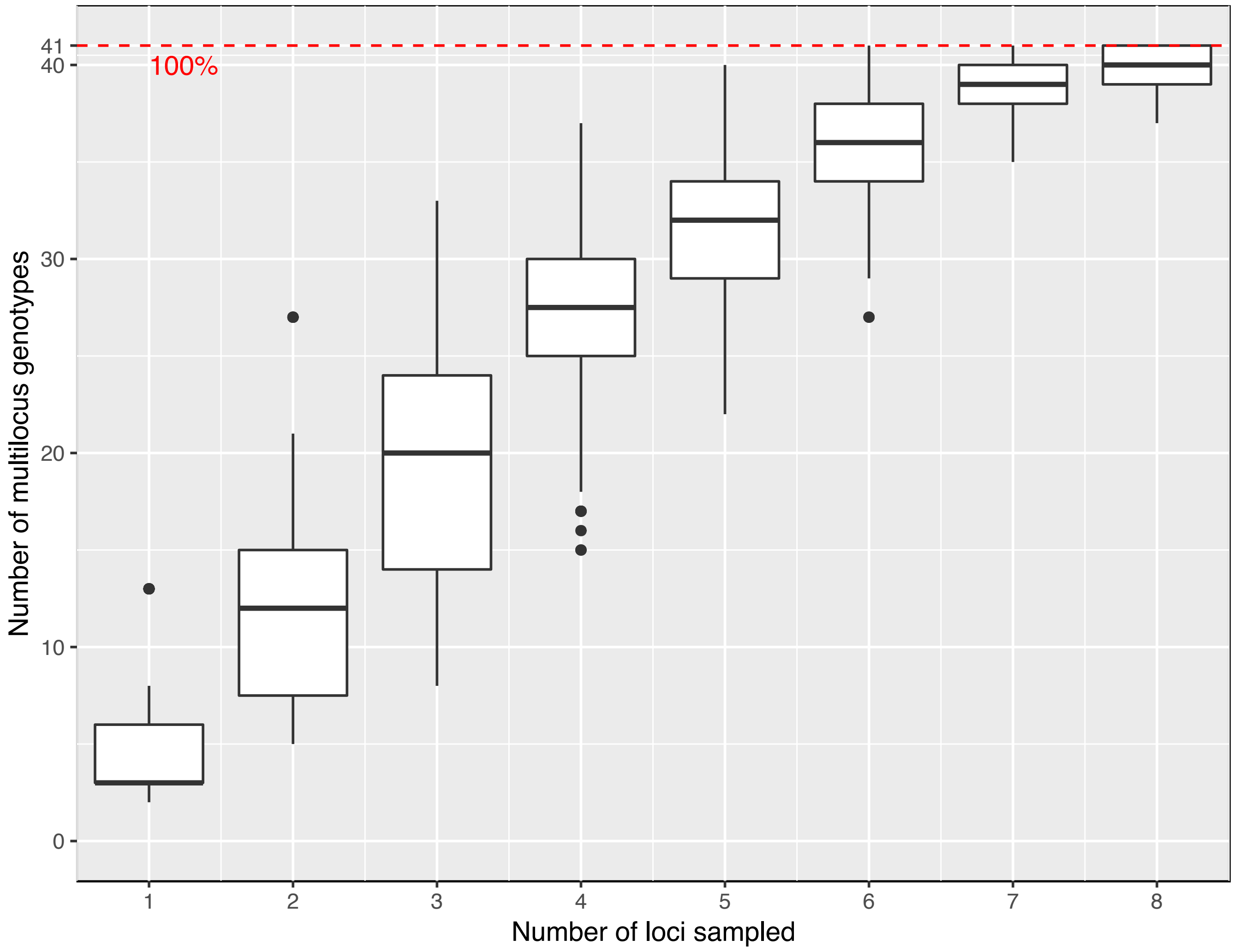


Figure S4: The relationship between number of microsatellite loci sampled and the total number of unique genotypes observed. Box plots indicate the medians, quartiles, and ranges of observed genotypes after randomly sampling 100 times. The dotted red line is the total number of genotypes (41) observed when measuring all nine loci.

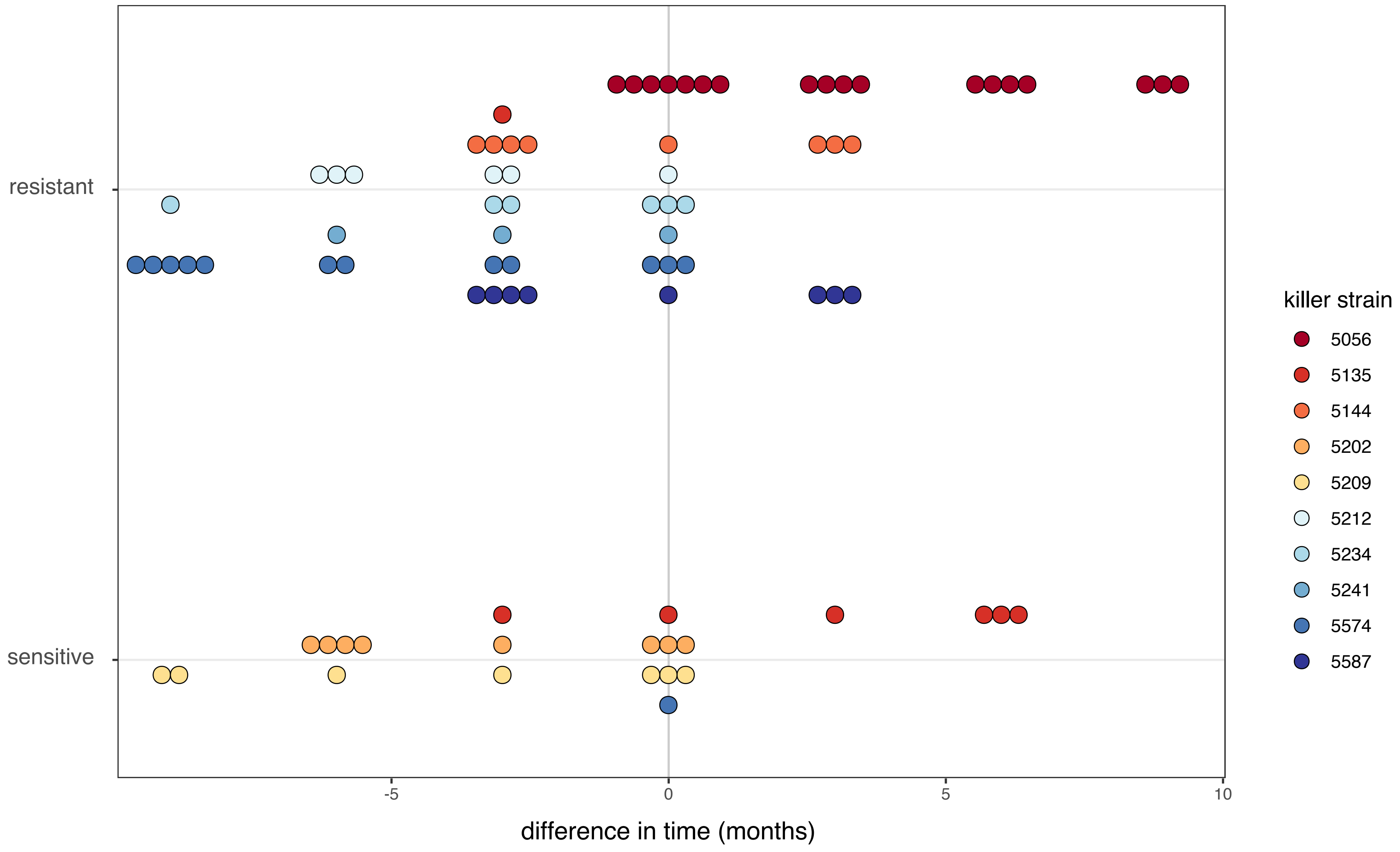


Figure S5: Resistance to killer toxins over time. Data are the same as for Figure 1c, but points are color-coded by the identity of the killer strain tested and sorted by whether resistance or sensitivity was observed. Each point represents a test of a single pair of killer and target strains.

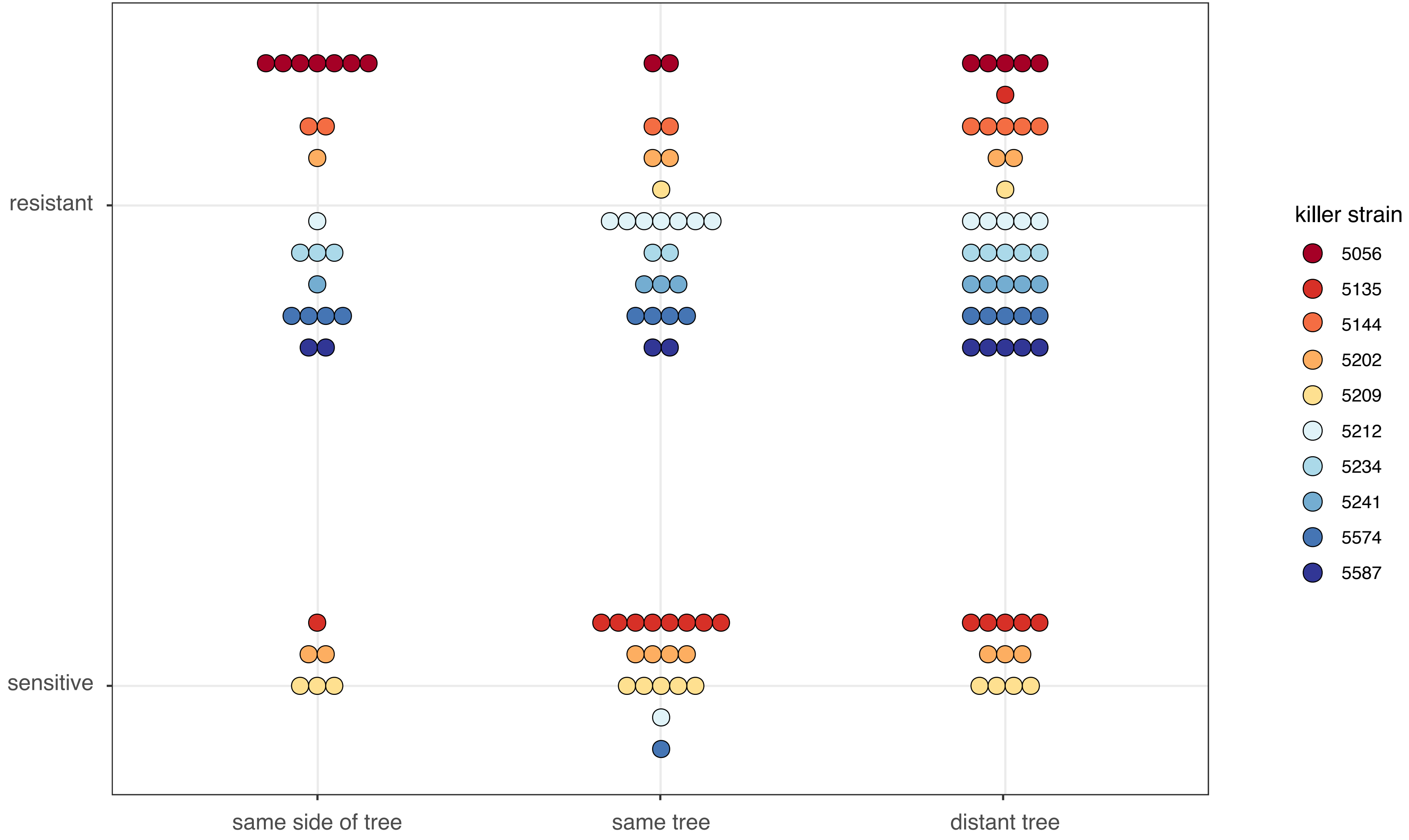


Figure S6: Resistance to killer toxins over space. Data are the same as for Figure 2c, but points are color-coded by the identity of the killer strain tested and sorted by whether resistance or sensitivity was observed. Each point represents a test of a single pair of killer and target strains.
